# Cellular cooperation shapes tumor growth: *a statistical mechanics mathematical model*

**DOI:** 10.1101/278614

**Authors:** Jeffrey West, Paul K. Newton

## Abstract

A tumor is made up of a heterogeneous collection of cell types all competing on a fitness landscape mediated by micro-environmental conditions that dictate their interactions. Despite the fact that much is known about cell signaling and cellular cooperation, the specifics of how the cell-to-cell coupling and the range over which this coupling acts affect the macroscopic tumor growth laws that govern total volume, mass, and carrying capacity remain poorly understood. We develop a statistical mechanics approach that focuses on the total number of possible states each cell can occupy, and show how different assumptions on correlations of these states gives rise to the many different macroscopic tumor growth laws used in the literature. Although it is widely understood that molecular and cellular heterogeneity within a tumor is a driver of growth, here we emphasize that focusing on the functional coupling of these states at the cellular level is what determines macroscopic growth characteristics.

**Significance statement:** A mathematical model relating tumor heterogeneity at the cellular level to tumor growth at the macroscopic level is described based on a statistical mechanics framework. The model takes into account the number of accessible states available to each cell as well as their long-range coupling (population cooperation) to other cells. We show that the degree to which cell populations cooperate determine the number of independent cell states, which in turn dictates the macroscopic (volumetric) growth law. It follows that targeting cell-to-cell interactions could be a way of mitigating and controlling tumor growth.

## 1. Introduction

A typical tumor is comprised of a remarkably heterogeneous agglomeration of cell types, both at the molecular level [1] and at the phenotypic/morphological level [2, 3]. Why this is often the case is still somewhat open to debate [4, 1], but clearly mutational instability [5, 6], ecological niches [7, 8], tissue micro-environmental factors [9, 10, 11] all contribute to this diversity of cell types which, in turn, enables natural selection to act to shape the fitness landscape of the tumor [3, 12], drive tumor growth [13, 14], and select for resistant sub-populations during treatment [15, 16]. A question we address in this note is what are the consequences of tumor heterogeneity at the cellular-scale with respect to key aspects of tumor growth at the macro-scale? It has long been known that cellular communication is essential for embryonic development, for example, but it has been less widely appreciated in cancer. In particular, how does cellular diversity, on the one hand, and cell-cell interactions and intercellular communication (via hormones,growth factors, neurotransmitters and cytokines mediated by gap junction channels) on the other hand, both work in concert to determine the macroscopic growth laws of a tumor? It has been observed that *the cancer problem is not merely a cell problem, it is a problem of cell interaction, not only within tissues, but with distant cells in other tissues* [17]. We know that communication processes among cells ultimately control many aspects of cell growth, including proliferation, differentiation, apoptosis, and a cell’s ability to adapt and respond to micro-environmental cues. We also know that disruption of cell-cell communication can lead to increased or decreased proliferation, abnormal differentiation, apoptotic alteration, and abnormal adaptive responses [18, 19, 20, 21]. In fact, it has been hypothesized that cancer essentially is a consequence of dysfunctional gap junctional intercellular communication [22, 23].

In addition, stochastic sub-cellular processes such as protein and mRNA expression are regulated by transcriptional, epigenetic, and signaling networks [24]. These complex interactions influence cell-cell variability within a population in non-intuitive ways that are often difficult to control experimentally, leading some to use statistical mechanics tools and informational theory methods to attempt to understand the molecular and cellular basis of stem cell pluripotency [25, 26]. An important information theory concept known as entropy has been proposed as a model of both inter-cell and intra-cell variability (a diversity scoring metric). One recent study analyzed single cell RNA-Seq profiles to calculate the entropy of the cellular transcriptome in context of the protein-protein interaction network to classify cell stemness [27]. Entropy, roughly a measure of the disorder of a population can be thought of as the probability governing the molecular state of a cell drawn from a population at random [28]. Thus, a population with a well-defined mRNA or protein expression will exhibit low entropy (low population variability or diversity) [26], whereas those with broadly diffused mRNA or protein expression profiles correspond to high entropy populations.

In this note, we frame the issue in terms of a statistical mechanics formulation based on the full range of available cellular micro-states to show how different short range and long range interaction rules shape tumor growth laws at the macroscopic level. Formulating population diversity and macroscopic growth into a statistical mechanics framework is useful in describing treatment implications as well. If an equilibrium distribution of cell states exists, to which every initial condition converges rapidly, the population is said to be *ergodic* [26]. Individual cells within an ergodic population can potentially independently explore the equilibrium state space, giving rise to an intrinsic robustness against targeted removal of specific population sub-types, a phenomenon observed experimentally with respect to a number of key pluripotent stem cell markers [29, 30]. Similar reconstructive populations have been experimentally observed in cancer cell phenotypic states, indicating the potential ergodicity of some cancers [18]. The collective action of all of these individual, coupled cellular interactions leads to some macroscopic behavior of the tumor at large. The purpose of this letter is to detail a framework for better understanding the collective behavior of inter-cellular coupling that shed light on these issues. In the next section we will link the diversity potential of the tumor with macroscopic growth laws, through the lens of statistical mechanics.

## 2. Heterogeneity, diversity, and growth

Genetic instability, a hallmark of cancer, is generally believed to be acquired early in tumorigenesis and thought to lead in a multi-step fashion to other cancer hallmarks [31, 32, 33]. This instability can be thought of as increasing the *potential* of diversity within a tumor leading to a large number of potential genetic or morphological ‘states’ each cell can occupy, which in turn gives rise to a combinatorial mushrooming of overall molecular/cellular configuration of the tumor. This makes a statistical mechanics approach to cancer modeling an attractive option. Kendal [34] introduced such a model based on these types of considerations which we take as a point of departure for our work, so it is useful to first review the main features of his simple argument. Consider a population of *n* cells where the *j*th cell has the potential to assume one of *q*_*j*_ possible states, *j* = 1, 2, 3, *…, n*. The number of combinations of states possible within the population is given as *P*, a measure of diversity potential of the tumor:

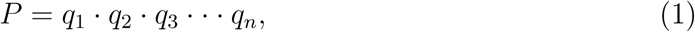

which can be related to the growth rate of the tumor. By introducing a function, *G*(*P*), that is a function of the number of proliferating cells, and using the requirements that *G* = *G*(*P*_1_ *P*_2_) = *G*(*P*_1_) + *G*(*P*_2_) for any two sub-populations *P*_1_ and *P*_2_ (additivity of *G*), we arrive at the requirement that *G* is logarithmic:

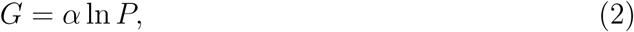

where ln is the natural logarithm. Then growth is proportional to *G*:

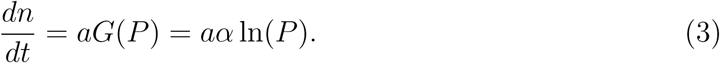

If we begin with the simplest completely uncoupled case where *q*_*j*_ = *m* (typically *m < < n*) for each *j* (i.e. each cell has available to it *m* possible states, with no coupling between the states), the combination of states is calculated from (1) as:

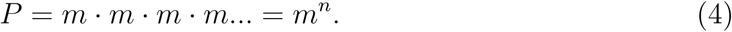

With this, we have the growth equation:

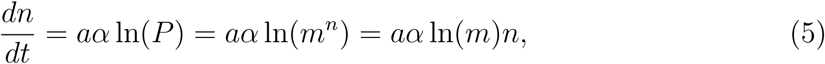

which yields exponential growth:

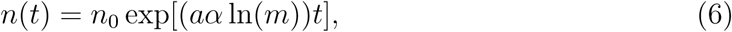

with growth rate proportional to the logarithm of the number of states available to each cell. We assume that *n*(*t* = 0) = *n*_0_, and that tumor volume *V* (*t*), in general is proportional to *n*(*t*). Therefore with no coupling among the cells to reduce the effective number of achievable states (degrees of freedom), tumor growth is effectively exponential. This is often the case in the very earliest stages of tumor growth before the cells have coordinated and synced.

By contrast, assume that the number of states each cell can achieve is reduced as the total cell population increases, due to functional coupling of the cell population. Instead of each cell acting independently as previously assumed, we now assume:

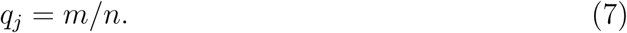

As *n* increases (i.e. as the tumor grows), the number of states each cell can achieve decreases linearly with *n*. With this assumption, we have:

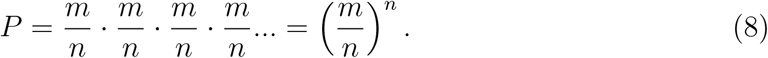

This gives rise to the growth equation:

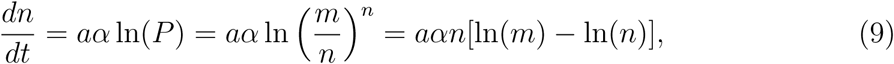

whose solution is the Gompertzian:

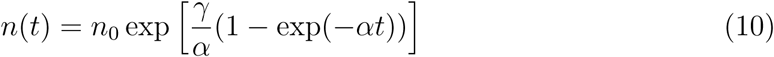

one of the most widely applied macroscopic tumor growth laws [35, 36, 37, 38, 39].

## 3. Cell population cooperation and volumetric growth

More generally, as the tumor grows, we assume that cell signaling and micro-environmental factors act to couple the states (much the way the uncoupled degrees of freedom in a mechanical system can sync, thereby reducing the effective degrees of freedom of the system as a whole). We write this as:

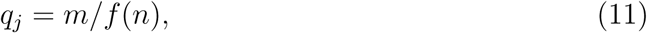

where the function *f* (*n*) determines the degree and details of the functional coupling among the states. Generally speaking, as the tumor grows, the functional coupling should increase (as in the Gompertzian case), hence *f* (*n*) should be an *increasing* function of *n*. With no loss of generality, we can write *f* (*n*) as the exponential of another function *g*(*n*) (to clean up the final equation):

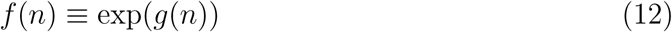

giving rise to the following relation:

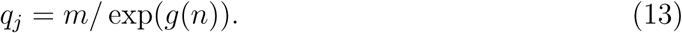

At this point, the functional form of *g*(*n*) is unspecified, but the denominator serves to restrict the total number of ‘free states’ *m*, a single cell can occupy. Using (1), the growth equation (2) then becomes:

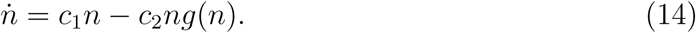

where *c*_1_ = *aα* ln(*m*), *c*_2_ = *aα*. The functional form *g*(*n*) determines which of the many common macroscopic growth models are in force (Exponential, Exponential-Linear, Logistic, Generalized Logistic, Gompertz, Von Bertanffly, and Power law). An overview of results are shown in Table 1. An example commentary on the use and history of common growth models in describing tumor dynamics can be found in Benzekry et al. [35].

**Table 1.**
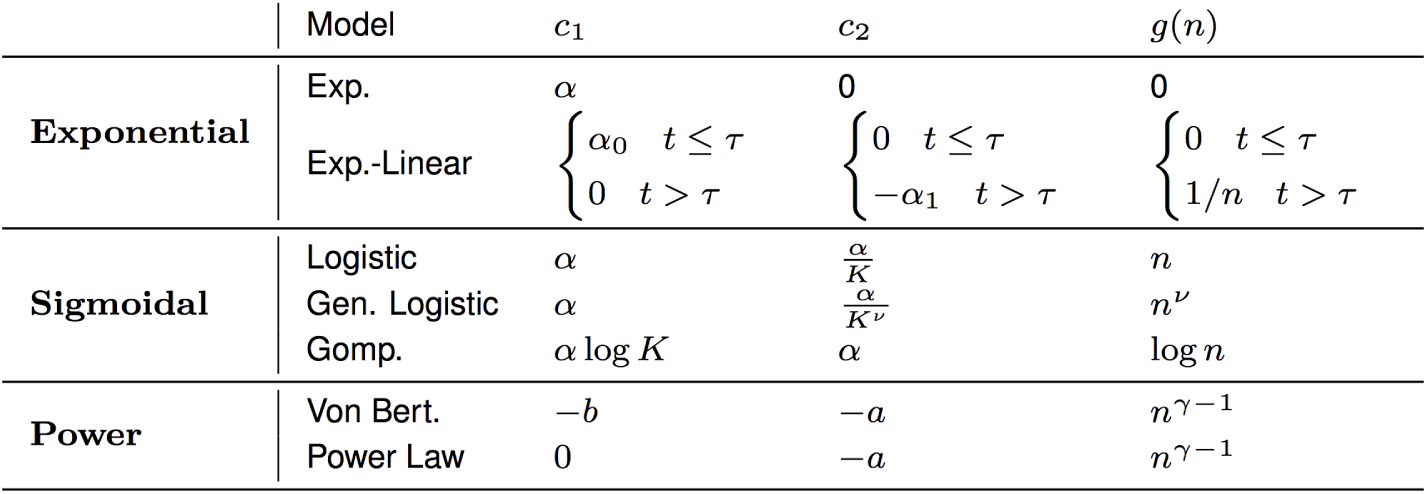
Overview of growth model parameters.

Broadly, the macroscopic growth models described below can be binned into three categories: 1) exponential, 2) sigmoidal, and 3) power law models. Exponential models are characterized by a long period of constant proliferation cell cycle time, while sigmoidal models have eventual slowed growth until an eventual plateau. A power law model also gives rise to a similarly slowed growth, but without a specified plateau. Each of these categories is associated with a certain functional coupling: approximately constant (exponential), increasing (sigmoidal), or decreasing (power). Explicit solutions for each of the growth models are detailed below, along with discussion on the implications of the form of coupling (i.e. the denominator of equation (13)).

## 4. Exponential Models

A simple model of tumor growth assumes a constant cell cycle time for all proliferating tumor cells, *T*_*c*_, which leads to exponential growth (eqns. (15),(16)). This model is also valid in the case of a constant fraction of the tumor volume is proliferating or that the cell cycle length is random variable with an exponential distribution (assuming independent, identically distributed individual cell cycles) [35].

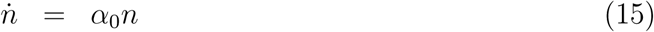

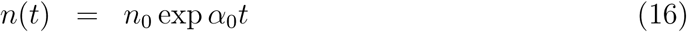

An extension of the exponential model assumes an initial exponential phase is followed by linear tumor growth (eqn. 17), first introduced here: [40].

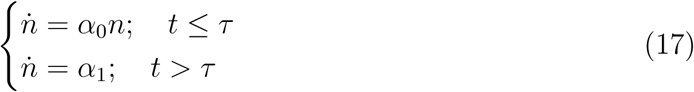

The exponential-linear model is solved explicitly below.

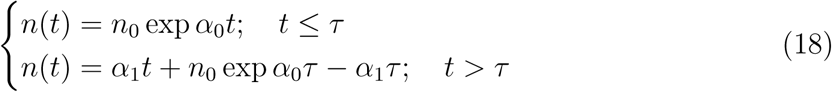

For the first phase of growth (*t < τ*), the coefficient *a*_0_ is equivalent to ln 2*/T*_*c*_, where *T*_*c*_ is the constant or exponentially distributed mean cell cycle time of the proliferative fraction within the tumor. Assuming a continuously differential function *τ* is uniquely determined as: 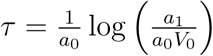 where *V*_0_ is the initial tumor volume [35, 40].

The exponential models are shown in figure 1a in blue (*α*_0_ = 1; *α*_1_ = 1*e*4; *τ* = 10). As explained above, a tumor grows from a single cell and the total number of ‘free states,’ m, a single cell can occupy is restricted by the exponential of a function form of *g*(*n*) (see equation 13). The forcing function determines how the coupling changes with an increasing tumor size, *n*. Exponential models are characterized by zero coupling (*g*(*n*) = 0; exponential), or approximately zero (*g*(*n*) *≈* 0; exponential-linear, for large *t*).

**Figure 1:**
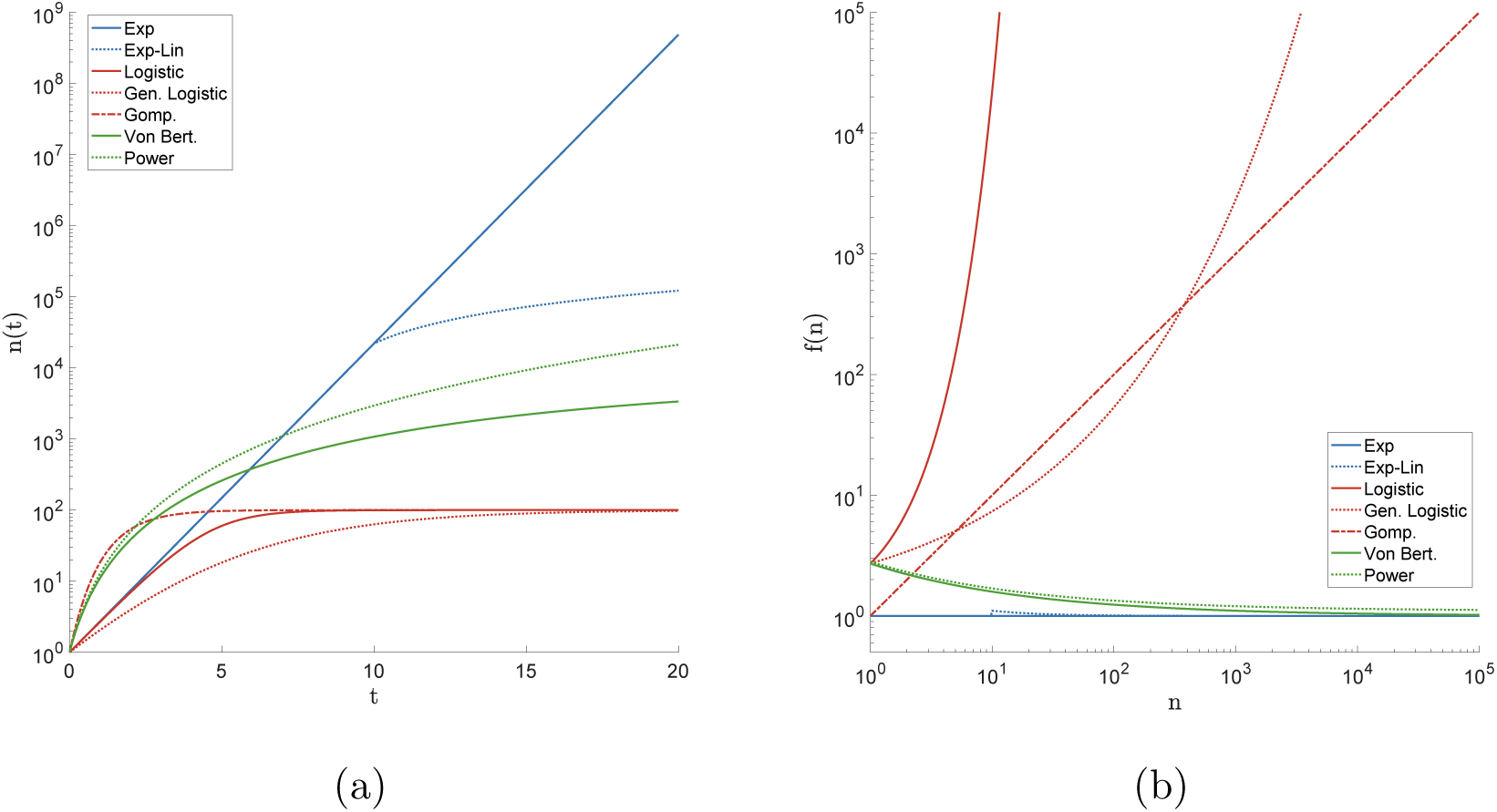
**(a)** Representative simulations of classical macroscopic models of tumor growth, shown here for *n*0 = 1 cell. Exponential models are shown in blue (*α*0 = 1; *α*1 = 1*e*4; *t* = 10). Sigmoidal models are shown in red (*α* = 1; *K* = 100; *ν* = 0.3). Power law models are shown in green (*γ* = 2*/*3; *a* = 4; *b* = 0.2); **(b)** Coupling function *f* (*n*) that determines the functional coupling of the cell states for identical parameters as referenced in figure 1a. There are three general classes of macroscopic growth models: exponential models (blue) characterized by (approximately) constant coupling; sigmoidal models (red) characterized by coupling that increases with tumor size; and power law models (green) characterized by coupling that decreases with tumor size.

## 5. Sigmoidal Growth models

The logistic (eqn. 19), generalized logistic (eqn. 21), and Gompertz (eqn. 24) equations are all in a general class of equations which quantify tumor growth in a sigmoidal shape where growth is slowed with increasing tumor size [35, 36, 41, 37].

Logistic growth (eqn. 19) is characterized by a linear decrease of relative growth rate and is often interpreted as a competition between proliferating tumor cells for space or nutrients. Logistic growth models have been used by many to describe tumor dynamics (e.g. [42, 37]).

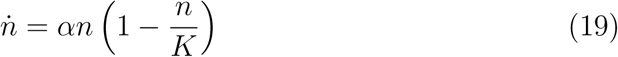

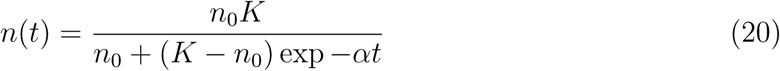

The logistic model can also be written as a generalized logistic equation, below.

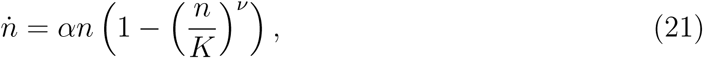

which is more conveniently written:

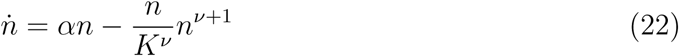

The explicit solution to the generalized logistic equation (eqn. 23) gives rise to the logistic model (eqn. 19) when *ν* = 1 [35].

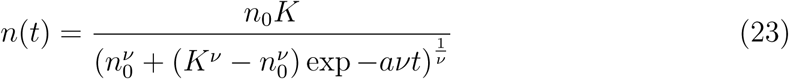

The generalized logistic model also converges to Gompertz model (see eqn. 24, below) when *ν →* 0 [35]. First introduced to describe human population growth [38], the Gompertz equation has also been used extensively in modeling tumor growth [36, 43, 39], Gompertzian growth is exponential growth with decaying growth rate:

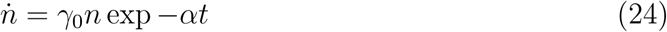

This is alternatively written:

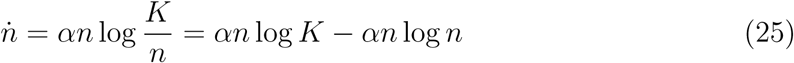

Where 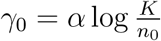. It is explicitly solved,

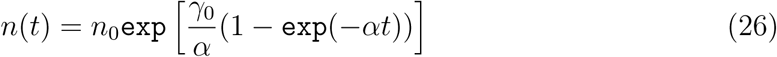

The sigmoidal models are shown in figure 1a in red (*K* = 100; *α* = 1; *ν* = 0.3;). The sigmoidal models are characterized by *increased* coupling (again, see denominator of equation 13) as the tumor size increases. This is shown in red in figure 1b and given by increasing function of *g*(*n*) in table 1. The forcing function for all sigmoidal functions are always increasing for values of *ν >*= 0.

## 6. Power Law Models

Another class of growth models, the power law model, has been used to derive general laws of tumor dynamics from a relationship between growth and metabolism [44, 45]. Metabolic rates within the tumor often scale with a power of the total tumor size [35, 44].

The Von Bertalanffy equation is written in eqn. 27, where the power law model is a special case, derived by neglecting the loss term (*b* = 0). Note: some often identify this model as the specific case *γ* = 2*/*3, termed “second type growth” [35]. Both the Von Bertalanffy model and the power law special case have been used to fit tumor spheroid data for murine models [46, 47] and breast cancer mammography screening data [48].

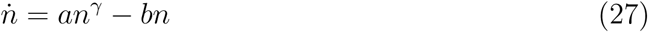

This model can be solved and written explicitly:

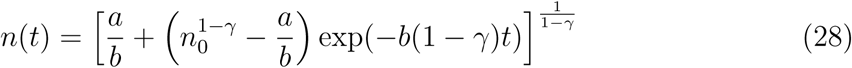

The power law models are shown in figure 1a in green (*γ* = 2*/*3; *a* = 4; *b* = 2). These models are characterized by *decreased* coupling as the tumor size increases. This is shown in green in figure 1b and given by decreasing functions of *g*(*n*) in table 1. Generally, the Von Bertalanffy model is associated with second type growth, with *γ* = 2*/*3, and the forcing function for all power functions are always decreasing for values of *γ <* 1. We now discuss the implications of the forcing functions in the three categories of models (exponential, sigmoidal, power).

## 7. Therapeutic implications

Traditional therapeutic treatments target the cancer cells directly by surgical removal or maximal eradication (chemotherapy and radiation). The linkage between cellular coupling and tumor growth leads naturally to the idea of therapeutic methods to try to disrupt or enhance the functional coupling of the cellular states in order to control tumor growth characteristics. This idea has been touched upon with the suggestion of multi-targeted therapeutic approaches disrupting or co-opting ecological interactions of tumor-host or tumor-tumor cell interactions. These approaches have been termed *ecological therapies* [21]. The model developed here provides a framework for determining how those interactions guide volumetric growth, which may lead to models that optimize timing of new therapies targeting cell-cell interactions, by targeting the mediators of those interactions. It is also important to note that tumor heterogeneity is not necessarily indicative of diversity potential. A highly heterogeneous tumor may also be a tightly coupled tumor. Increased heterogeneity may lead to many other evolutionary advantages (for example, fostering the selection of a resistant subclone [31]), but limits possible future cell states, which slows tumor growth.

In general terms, the growth law that a tumor follows has definitive therapeutic implications. Cancers that follow exponential growth laws can be assumed to have constant cell-cell interactions (or lack thereof, in the case of blood cancers) and targeted therapies may have more promise in these scenarios, while the therapies aimed at limiting cell-cell interactions described previously will likely have little effect. Likewise, growth models with decreased coupling over time (green curves in figure 1b) will benefit from ecological therapies early in tumor development but not as tumor sizes increase to clinical relevant sizes. Otherwise, growth laws associated with increased coupling (red curves in figure 1b) show promise with respect to therapies targeting interactions.

Some have described the therapeutic implications of tumors viewed as ecosystems, characterized by the numbers of each type of clone/subclone, their spatial and temporal assortment, and their interactions with each other and their physical and chemical microenvironments [21, 49]. Much research has been done in the area of collateral sensitivity, to determine if there is pharmacological interaction (additivity or synergy) of multiple drugs in combination or sequence (see [50, 51, 52]), showing that drugs may have complex downstream effects on cell-cell interactions. Communication or feedback between tumor cells may provide negative (competition, predation, amensalism, parasitism) or positive (commensalism, synergism, mutualism) [21, 20] growth. Organisms compete for limited resources and cooperate for mutual advantage with interactions fluctuating with resource consumption or cell turnover. Additionally, tumor cell interactions may also be dependent on benefits derived from non-cancer cells (endothelial cells, cancer-associated fibroblasts, and tumor-associated macrophages) [21, 20]. Targeting these non-cancer cells, from which the cancer cells are receiving benefit, should also provide therapeutic benefits during the process by which a tumor transitions from a closed system (primary tumor) to an open system of circulating cells to distant colonies (metastatic cancers) – this too has implications on resource limitation and cell-cell interactions. But a clear understanding of the relationship between cell cooperation and volumetric growth is a necessary step in the direction of exploiting these connections.

